# Genotype prediction of 336,463 samples from public expression data

**DOI:** 10.1101/2023.10.21.562237

**Authors:** Afrooz Razi, Christopher C. Lo, Siruo Wang, Jeffrey T. Leek, Kasper D. Hansen

**Author notes:** Correspondence to (KDH), (JTL). Equal contributions as first authors (Afrooz Razi, Christopher C. Lo).

## Abstract

Tens of thousands of RNA-sequencing experiments comprising hundreds of thousands of individual samples have now been performed. These data represent a broad range of experimental conditions, sequencing technologies, and hypotheses under study. The Recount project has aggregated and uniformly processed hundreds of thousands of publicly available RNA-seq samples. Most of these samples only include RNA expression measurements; genotype data for these same samples would enable a wide range of analyses including variant prioritization, eQTL analysis, and studies of allele specific expression. Here, we developed a statistical model based on the existing reference and alternative read counts from the RNA-seq experiments available through Recount3 to predict genotypes at autosomal biallelic loci in coding regions. We demonstrate the accuracy of our model using large-scale studies that measured both gene expression and genotype genome-wide. We show that our predictive model is highly accurate with 99.5% overall accuracy, 99.6% major allele accuracy, and 90.4% minor allele accuracy. Our model is robust to tissue and study effects, provided the coverage is high enough. We applied this model to genotype all the samples in Recount3 and provide the largest ready-to-use expression repository containing genotype information. We illustrate that the predicted genotype from RNA-seq data is sufficient to unravel the underlying population structure of samples in Recount3 using Principal Component Analysis.

## Background

RNA sequencing technologies are now commonplace for collecting transcriptome data (Mortazavi et al., 2008). As technologies have improved and costs have dropped, high-throughput expression data has accumulated quickly in public archives (Langmead, Nellore, 2018). Initiatives on providing public access to available expression data has promoted reproducibility and re-usability of the existing data (Kaminuma et al., 2010; Benson et al., 2012; Leinonen et al., 2011; Mailman et al., 2007). However, there are challenges in using these population scale genomic data. To address this, we previously developed the Recount3 repository, which provides public access to uniformly processed expression summaries aggregated from raw RNA sequencing data available as part of the unrestricted access data from SRA, and dbGAP protected data from Genotype Tissue Expression (GTEx version 8), and The Cancer Genome Atlas (TCGA) (Wilks et al., 2021; Collado-Torres et al., 2017). By doing so, Recount3 reduces the inevitable computational costs involved with preprocessing and analyzing large-scale public gene expression data and makes them more accessible to researchers. We have also developed machine learning approaches to predict metadata - including tissue type, sex, and other technological variables - to improve the value of this large collection of integrated transcriptomic data sets (Ellis et al., 2018).

While these gene expression data are valuable on their own, many questions in functional genomics can only be answered if both gene expression and genotype data are available on the same samples. Unfortunately, for most gene expression samples in Recount3 we do not have corresponding information on genetic variants. Important exceptions are GTEx, TCGA and the Geuvadis project (TG Consortium et al., 2013).

Here, we aim to extend Recount3 repository by predicting sample genotype directly from the RNA sequencing summaries calculated by the Recount3 project. De-novo variant calling from RNA-seq data has gained popularity in non-human studies due to lower costs and lack of an accurate reference genome (Adetunji et al., 2019; Jehl et al., 2021; Bakhtiarizadeh, Alamouti, 2020). Genotype calling based on the reference genome has been widely used for human samples using the Genome Analysis Toolkit (GATK) (DePristo et al., 2011; Long et al., 2022; Tang et al., 2014; Deelen et al., 2015; Vigorito et al., 2023). However, the GATK tool introduces two problems for genotyping in Recount3; 1) To overcome the storage burden, Recount3 does not provide access to raw FASTQ files which is needed as an input for GATK pipeline. 2) GATK read alignment and quality control steps are redundant in Recount3 genotyping since the samples are already uniformly processed. Instead, we sought to use the compressed gene expression summaries made available as part of Recount3 to reduce computational complexity, storage costs, and algorithmic costs for genotype prediction.

Here, we develop a machine learning model to genotype samples from RNA-seq counts in Recount3 human samples. Our approach uses the reference and alternate allele counts for biallelic SNPs in the coding regions that are produced by Recount3. For each genotype, we model the relationship between transformed alternate allele count and transformed reference coverage. Our approach allows single-sample processing. We estimate out-of-sample, within-study accuracy using a held out GTEx test set and out-of-study accuracy using data from the Geuvadis project. To show that our predicted genotypes contains biological signal of population structure, we developed a method to assign 1000 genomes super-population groups to both bulk and single cell RNA-seq samples; the largest analysis of inferred genetic ancestry to public expression data. In bulk RNA-seq data, we have 35.9% European samples and 38% Admixed Americans. This parallels results from genetic sequencing data which has been shown to be biased toward Europeans (Popejoy, Fullerton, 2016; Sirugo et al., 2019).

We applied our genotyping model to predict genotypes for 336,463 human samples in Recount3. We have developed the RecountGenotyper R package that can be used to genotype future samples. The resulting genotypes enable studies of the relationship between gene expression and genetic variation at an unprecedented scale.

## Results

We developed a model to genotype RNA-seq samples using gene expression summaries available through the Recount3 project. Recount3 uses a customized workflow to process a large amount of data and as a consequence, BAM files containing read alignments are discarded during processing. This prohibits the application of standard workflows for genotyping RNA-seq samples. Instead, the available data on genotypes are in the form of so-called alternative counts files. For each base in the genome, these files record the coverage of each of the alternative nucleotides compared to the reference genome (GRCh38) (Figure 1). We generated alternative counts files as part of the Recount3 processing, but these files are not publicly available to avoid sample re-identification.

**Figure 1.**
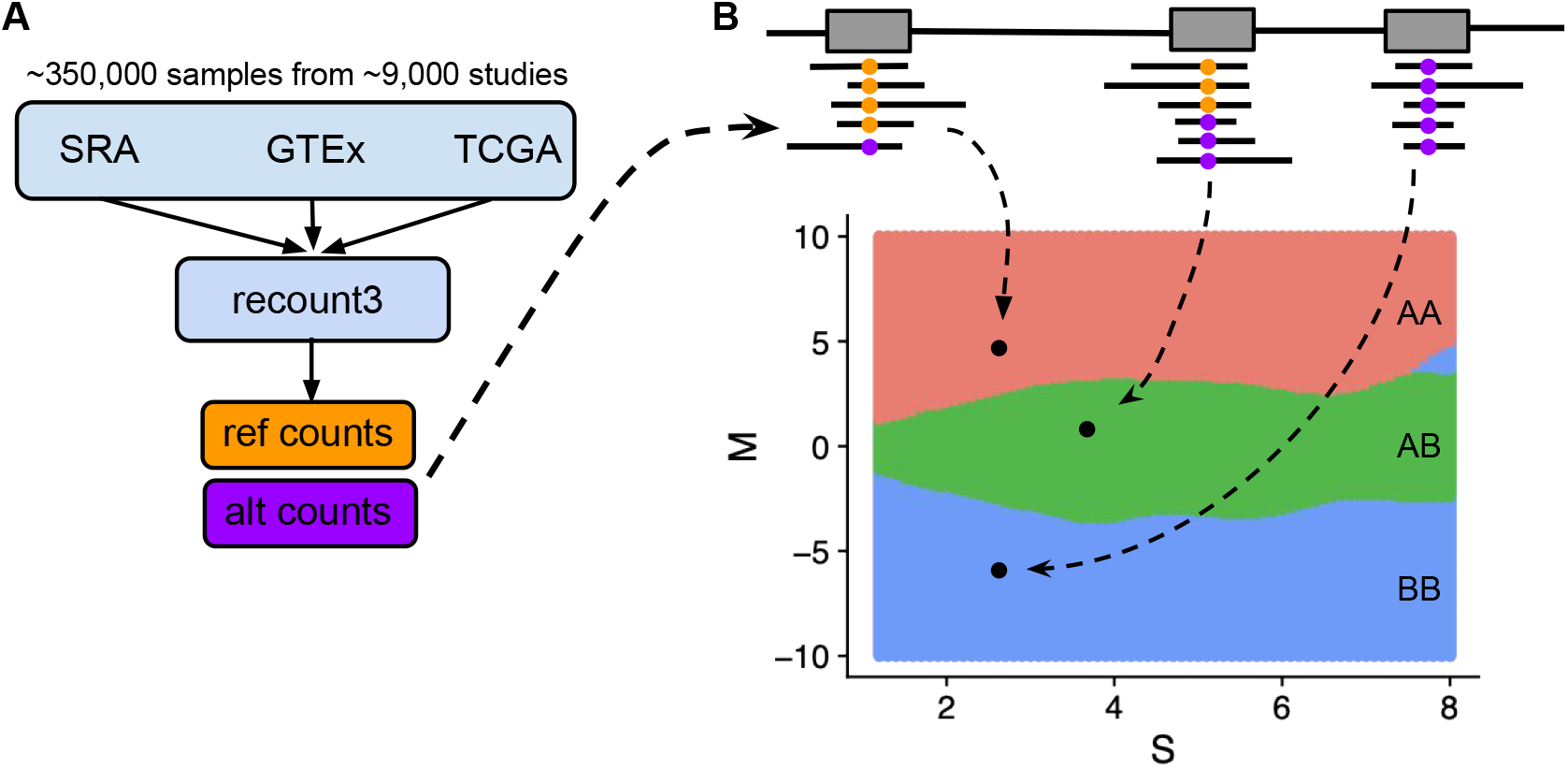
Genotyping workflow for genotype calling from Recount3 RNA-seq data. **(A)** Recount3 includes processed RNA-seq data from about 350,000 human samples aggregated from SRA, GTEx, and TCGA. Gene expression summaries are in the form of total read counts and alternative read counts. We subtract total read counts from alternative read counts to get reference counts. **(B)** Given a SNP from a sample to be genotyped, we transform reference and alternative read counts to M and S values. Based on M and S values, we predict our genotype using the MAP estimate. The decision boundaries for the MAP estimate are colored in red, green, and blue for reference homozygous genotype (AA), heterozygous (AB), and alternative homozygous (BB), respectively.

While Recount3 contains samples from both human and mouse, our focus here is on human samples. Using the methods described below, we successfully genotyped 316,639 human samples originating from unrestricted access SRA data, 19,081 samples from GTEx, and 743 non-cancer samples from TCGA. The remaining 10,630 cancer samples from TCGA were excluded due to potential chromosomal abnormalities that might affect allele read counts. We will refer to each subset of Recount3 data as SRA, GTEx, and TCGA throughout the paper unless stated otherwise.

To reduce our genotyping error rate, we only genotype samples on a pre-specified set of locations of known single nucleotide polymorphisms (SNPs). First, this set of SNPs only include autosomal biallelic SNPs in human exons. Second, because we used the GTEx data for model training, we furthermore restricted the set of SNPs to variants which are polymorphic across the 838 individuals with whole-genome DNA sequencing data in GTEx version 8. This resulted in an initial list of 20,980,266 SNP locations (Methods).

### Genotyping model and prediction

The input data to our model is the coverage of the reference and the alternative alleles for each of the candidate SNPs (Methods). The reference allele is defined as the allele present in the reference genome used for alignment (GRCh38). We next transformed the coverage of the reference and alternative allele into *M* and *S* values (Supplemetary Figure S1), for SNP *i* from a sample.

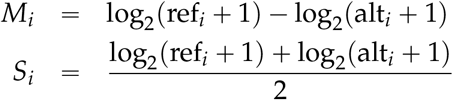

We then removed candidate SNPs with an average coverage across the training set of less than 3. A scatterplot of M vs. S reveals 3 distinct clusters representing 3 genotypes (Figure 2a).

**Figure 2.**
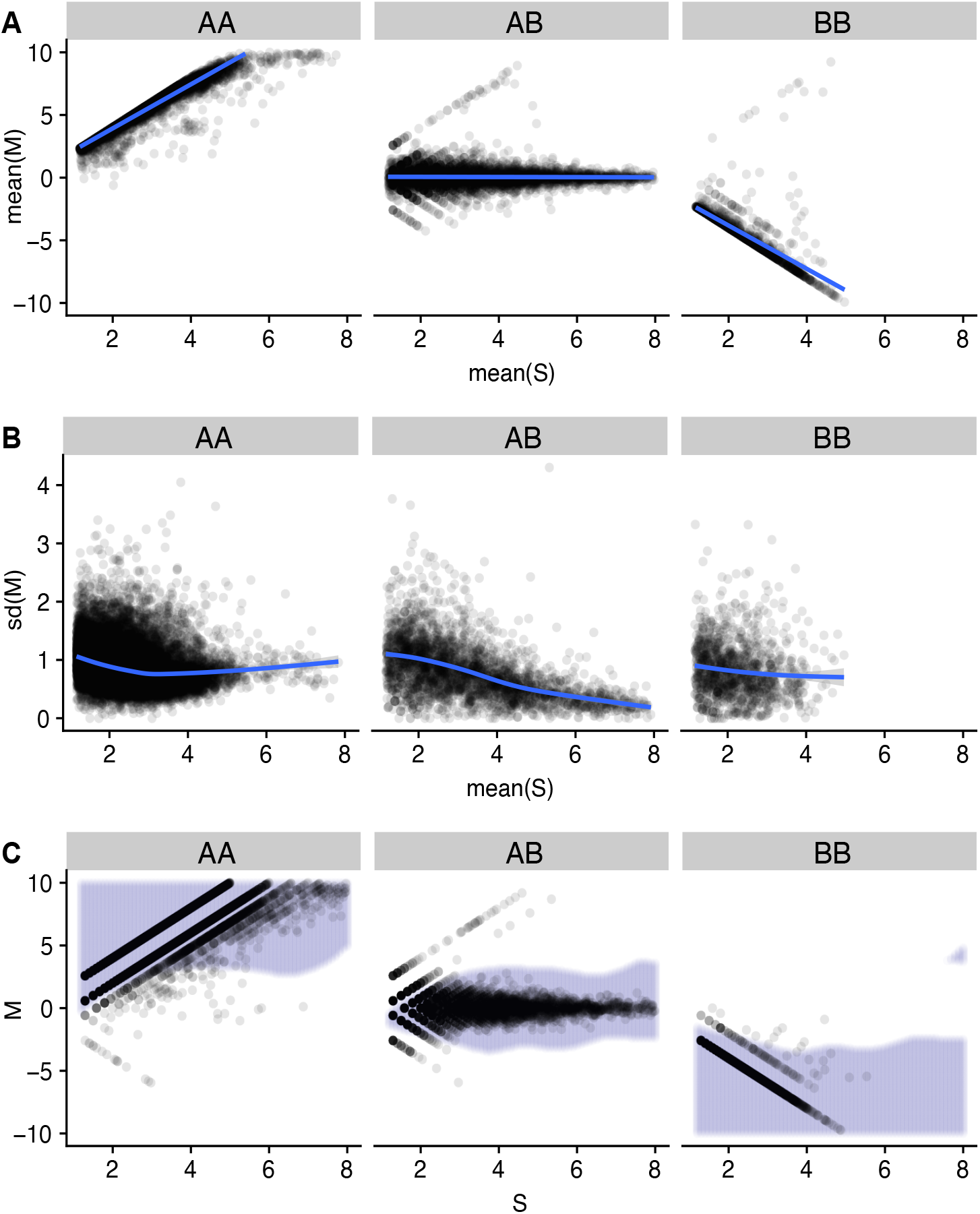
Model development on GTEx training samples. 200 samples were randomly selected from the GTEx training set to illustrate the model building process. Each plot is faceted based on AA (homozygous reference) genotype, AB (heterozygous), and BB (homozgyous alternative). **(A)** Each point represents a SNP’s mean M and mean S values across training samples, with a linear fit describing the relationship. **(B)** Each point represent a SNP’s standard deviation of M and mean S values across training samples, with a GAM fit describing the relationship. **(C)** Each point represent a SNP’s M and S values from a single sample. The decision boundary of the fitted model on the training set is colored in lilac.

We used this representation to build our model, which we name RecountGenotyper. Genotype was modeled as a latent discrete variable with 3 genotype states (AA, AB, BB), with “A” always referring to the reference allele and associated with positive values of *M*. Based on Figure 2, we assume a linear relationship between *M* and *S* conditional on genotype, with a flexible specification of the variance of *M* conditional on *S* (Methods). We use the maximum a posteriori probability (MAP) estimator to predict genotypes as a function of *M* and *S*; this results in the prediction boundaries depicted in Figures 1B, 2C. Our model represents a relationship between *M* and *S* learned jointly across all training SNPs.

For prediction on a single sample, we first select the subset of candidate SNP locations which has a sample-specific coverage greater than 4. We then use sample-specific *M* and *S* values to predict genotypes for these specific SNPs. Sensitivity analysis on the prior distributions of genotypes suggests that the choice of prior has negligible impact on the prediction results (Supplementary Figure S2). In practice, we use the genotype distribution in GTEx as the prior. After the model has been trained, this approach allows for single-sample processing.

For model training we use RNA-seq samples from the GTEx consortium version 8. We used samples from individuals which have been genotyped using whole-genome DNA sequencing and we use these genotype calls as gold-standard for model training. We created a training set by selecting samples from 638 individuals and 33 tissues. Each tissue type is balanced in the training set to account for variability of tissue expression. The remaining 200 individuals were used as the test set which consists of 3901 samples across 54 tissues (Methods). There was no overlap in the individuals in the training and test set.

### Evaluating model performance

The mean overall allelic accuracy for the GTEx test set of 16,185,565 unique SNPs is 99.5% (Figure 3A). We genotyped 3,901 samples from 200 individuals in our GTEx test set and computed the SNP accuracy for all samples using whole-genome DNA sequencing genotypes as the gold-standard. The SNP accuracy can be reported at either the genotype or the allelic level, and we used allelic accuracy to evaluate model performance. The advantage of allelic accuracy is that a prediction of AB is considered better than a prediction of BB when the true genotype is AA. Because we are predicting biallelic SNPs, in each sample each SNP will have an accuracy of either 0/2 = 0, 1/2 = 0.5, or 2/2 = 1.

**Figure 3.**
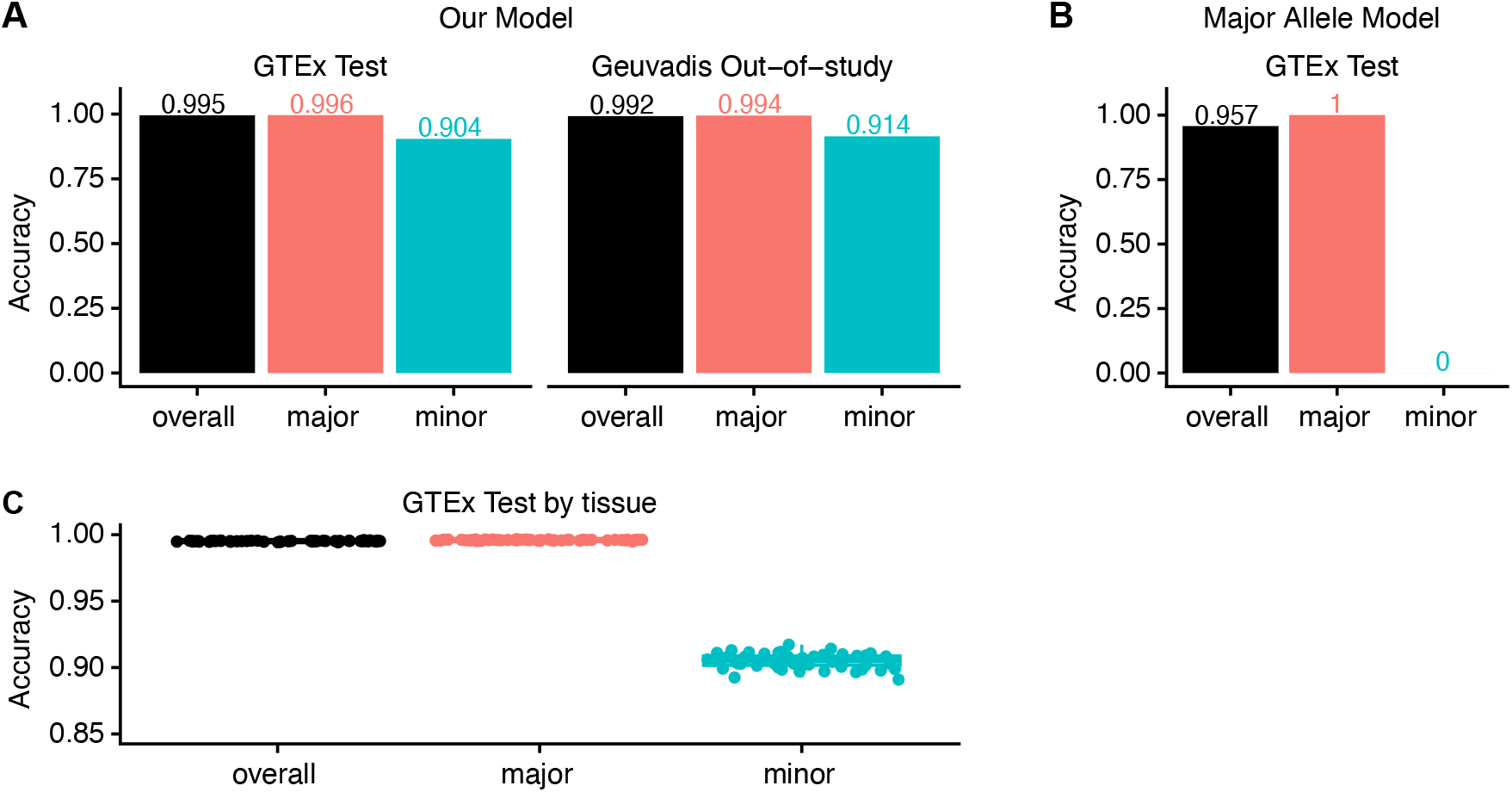
Evaluating model performance using overall and conditional allelic accuracy. (Conditional) allelic accuracy for GTEx (3,091 samples from 200 individuals) and Geuvadis test sets (462 individuals). **(A)** Model performance on GTEx and Geuvadis test sets. **(B)** The performance of the major allele model, always predicting the major allele. **(C)** Model performance across 54 tissues in GTEx.

We reasoned that the overall accuracy would be strongly affected by the major allele frequency distribution in the test set, where the mean major allele frequency is 96.6% (Supplementary Figure S3). To illustrate this, we evaluated overall accuracy for a model which always predicts the major allele across the 838 GTEx individuals, termed the “major allele model”. This model achieves a high overall accuracy of 95.7% (Figure 3B). To parse out the dominating effect of the major allele on the overall accuracy when the mean major allele frequency is high, we additionally report our accuracy conditional on the major and minor allele (Methods, Table 1). We define major and minor allele based on 838 individuals in GTEx version 8. On the GTEx test set, our model achieves a mean major allele accuracy of 99.6%, and a mean minor allele accuracy of 90.4% (Figure 3A). For comparison, the major allele model achieves a mean major allele accuracy of 100% and mean minor allele accuracy of 0% (Figure 3B).

**Table 1.**
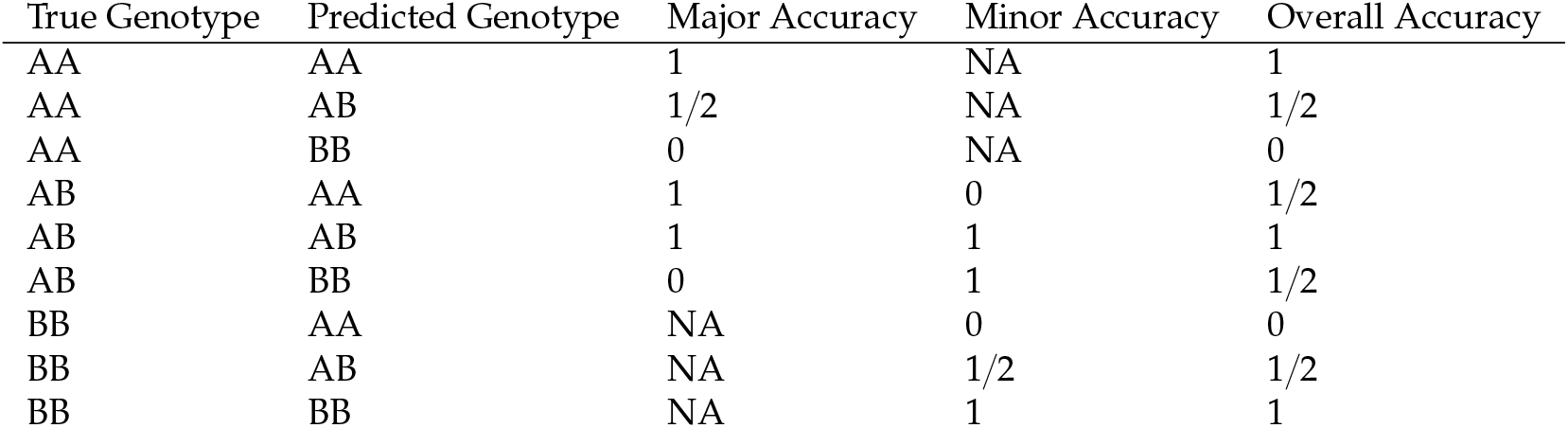
Rule for computing genotyping accuracy at the allele level. The major allele is denoted as “A”, and the minor allele is denoted as “B”. Given the true genotypes, the major, minor, and overall accuracy can be calculted for each SNP.

| True Genotype | Predicted Genotype | Major Accuracy | Minor Accuracy | Overall Accuracy |
| --- | --- | --- | --- | --- |
| AA | AA | 1 | NA | 1 |
| AA | AB | 1/2 | NA | 1/2 |
| AA | BB | 0 | NA | 0 |
| AB | AA | 1 | 0 | 1/2 |
| AB | AB | 1 | 1 | 1 |
| AB | BB | 0 | 1 | 1/2 |
| BB | AA | NA | 0 | 0 |
| BB | AB | NA | 1/2 | 1/2 |
| BB | BB | NA | 1 | 1 |

To ensure that our model was not overfitted to the GTEx study, we used the Geuvadis project as an out-of-study test set (TG Consortium et al., 2013). We computed the accuracy of 1,373,457 expressed SNPs from 462 Lymphoblastiod Cell Lines (LCL) in Geuvadis, using genotypes imputed from SNP arrays as the gold-standard. We attained a mean overall allelic accuracy of 99.2%, mean major allelic accuracy of 99.4%, and mean minor allelic accuracy of 91.4%, which is on par with the GTEx test set performance (Figure 3A). Note that the major allele is defined using GTEx individuals, despite the performance measure being computed on Geuvadis samples; this may explain the higher minor allelic accuracy for Geuvadis compared to the GTEx test set. We additionally compared the Geuvadis result to GTEx lymphoblastoid cell line test samples, and observe very similar results (Supplementary Figure S4).

Our model performance is robust to variations in tissue expression patterns. Previous studies of tissue specific gene expression have shown that samples from the same tissue type tend to express a common set of genes unique to the tissue (Lonsdale et al., 2013). Since our model only predicts SNP genotype at expressed genes, we expect to genotype a similar set of SNPs for samples from the same tissue type. In contrast, we expect to see variation in the set of SNPs genotyped when looking across tissues. Thus, we investigated the model performance for each tissue type. Accuracy for each tissue in the GTEx test set was evaluated on an average of 4,097,750 expressed SNPs (min: 982,965 SNPs, max: 7,478,944 SNPs) and an average of 72 samples (min: 1 sample, max: 182 samples). We see near-uniform mean overall accuracy across tissues with the lowest of 99.4% and highest of 99.6%, near-uniform mean major accuracy with the lowest of 99.5% and highest of 99.7%, and very uniform performance of mean minor allele accuracy with the lowest of 89.0% and highest of 91.7% (Figure 3C).

### Modelling genotyping accuracy

To help user interpretation of our predicted genotypes, we sought to develop a model for genotyping accuracy when the gold-standard genotype is unknown. This makes it possible to further select predicted genotypes based on their expected accuracy. We shift our focus from allelic accuracy to genotyping accuracy as we believe the latter to be more important to the end user.

To develop our model, we observe that genotyping accuracy is highly dependent on sequencing coverage and major allele frequency. Using 1,056,408 SNPs from 638 samples in the GTEx training set, we computed mean genotyping accuracy of SNPs that have the same coverage and similar range of major allele frequency. We observe that mean genotyping accuracy increases as a function of sequencing coverage; the rate of change is dependent on the major allele frequency (Supplementary Figure S5). Therefore, we model genotyping accuracy using a logistic regression model where we use a smooth function of coverage interacting with 6 discrete major allele frequency bins (Methods).

To evaluate model performance, we use the GTEx test set and the Geuvadis out-of-study test test, and observe excellent performance (Figure 4). For the GTEx test set, the mean absolute error between predicted and observed mean genotyping accuracy is 0.5% (min: 0%, max: 5.1%) for sequencing coverage ⩽ 30. For the Geuvadis out-of-study set, the mean absolute error is also 0.5%. Two tissues – whole blood and LCLs from GTEx – perform less well: the model over-predicts the true genotyping accuracy (Supplementary Figure S6). However, the mean absolute error remains small, with 1.3% for GTEx whole blood and 1.5% for GTEx LCLs.

**Figure 4.**
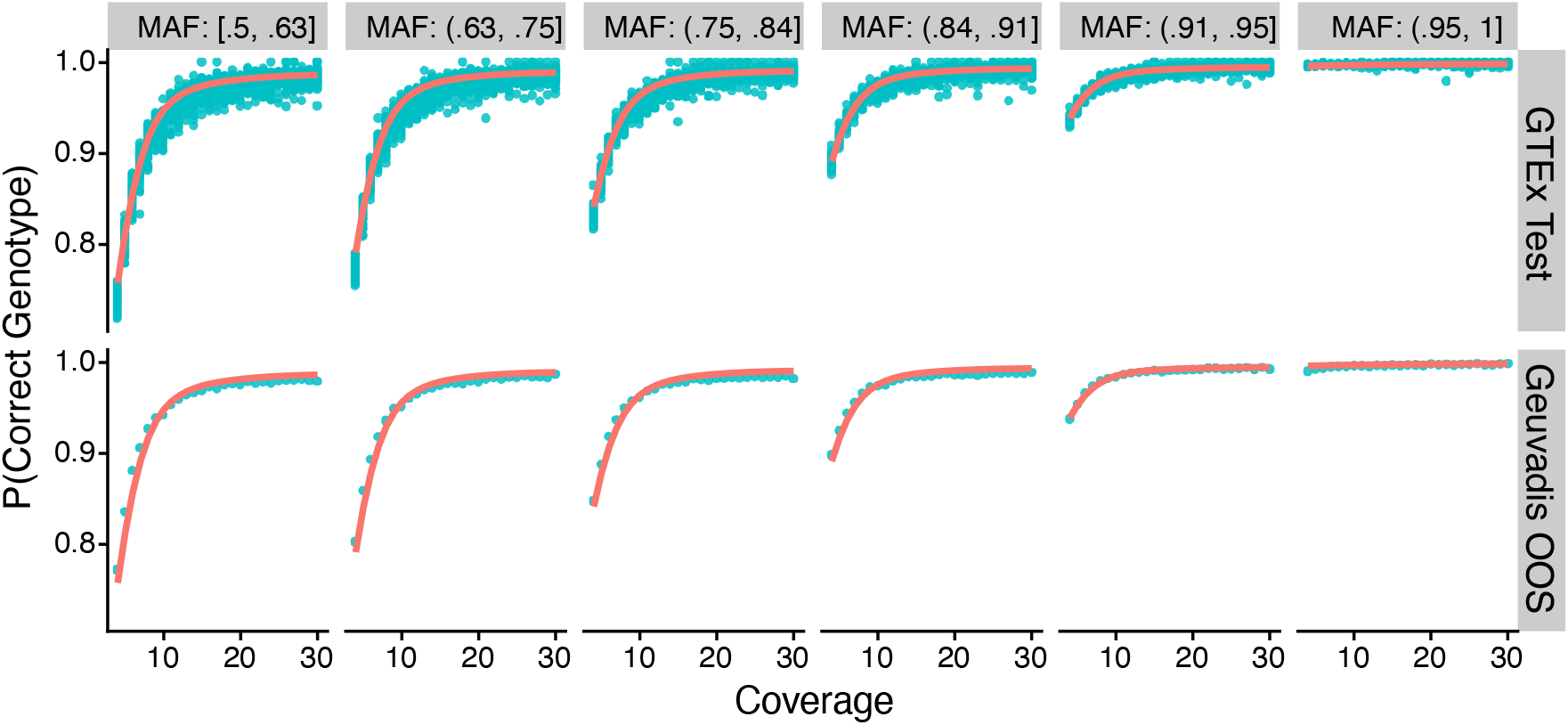
Accuracy prediction model on GTEx test set and Geuvadis out-of-study. Observed genotyping accuracy at the SNP level is grouped by discrete coverage values and major allele frequency bin to get a continuous averaged accuracy value (blue-green points). The model prediction is shown in red lines. Each facet along the x-axis denotes a major allele frequency bin. Each facet along the y-axis is the test set name.

### Genotyping SRA samples

After our promising genotype and accuracy model performance, we applied our model to the entire 315,443 SRA human samples available in Recount3. A substantial amount of these samples are single-cell (Wilks et al., 2021). We successfully genotyped an average of 932,460 SNPs per bulk and 90,863 per single-cell RNA-seq samples. On average, 1% SNPs in a bulk-RNA seq sample from SRA have coverage > = 15 to be genotyped with predicted near-perfect accuracy. For single-cell RNA-seq samples, 0.2% SNPs have coverage > = 15.

### Population structure analysis of predicted genotypes

The representation of diverse populations in public genetics data is of importance to precision medicine (Popejoy, Fullerton, 2016; Petrovski, Goldstein, 2016). Most (91%) genome-wide association studies on complex traits have been performed on European ancestry populations (Fitipaldi, Franks, 2023). This raises the question of the population composition of publicly available RNA-seq data (Barral-Arca et al., 2019; Harismendy et al., 2019). Existing approaches to inferring population structure from RNA-seq data consists of performing principal component analysis of genotypes from a select panel of SNPs; such an unsupervised analysis requires identifying which principal component represents which populations (Fachrul et al., 2023).

To unravel the population structure of samples in Recount3, we projected our predicted genotypes from RNA-seq data onto a reference subspace that describes the population diversity of the 1000 Genomes Project, which consists of a diverse set of 2,504 individuals from 26 populations grouped into 5 super-populations. We generated the reference population subspace by performing principal component analysis (PCA) on whole-genome DNA sequencing data from these 2,504 individuals. As expected, the first two principal components separate the 5 super-populations (Figure 5A). We scaled the axes based on the variance explained by each PC as recommended by best practices for dimensional reduction analysis (Nguyen, Holmes, 2019).

**Figure 5.**
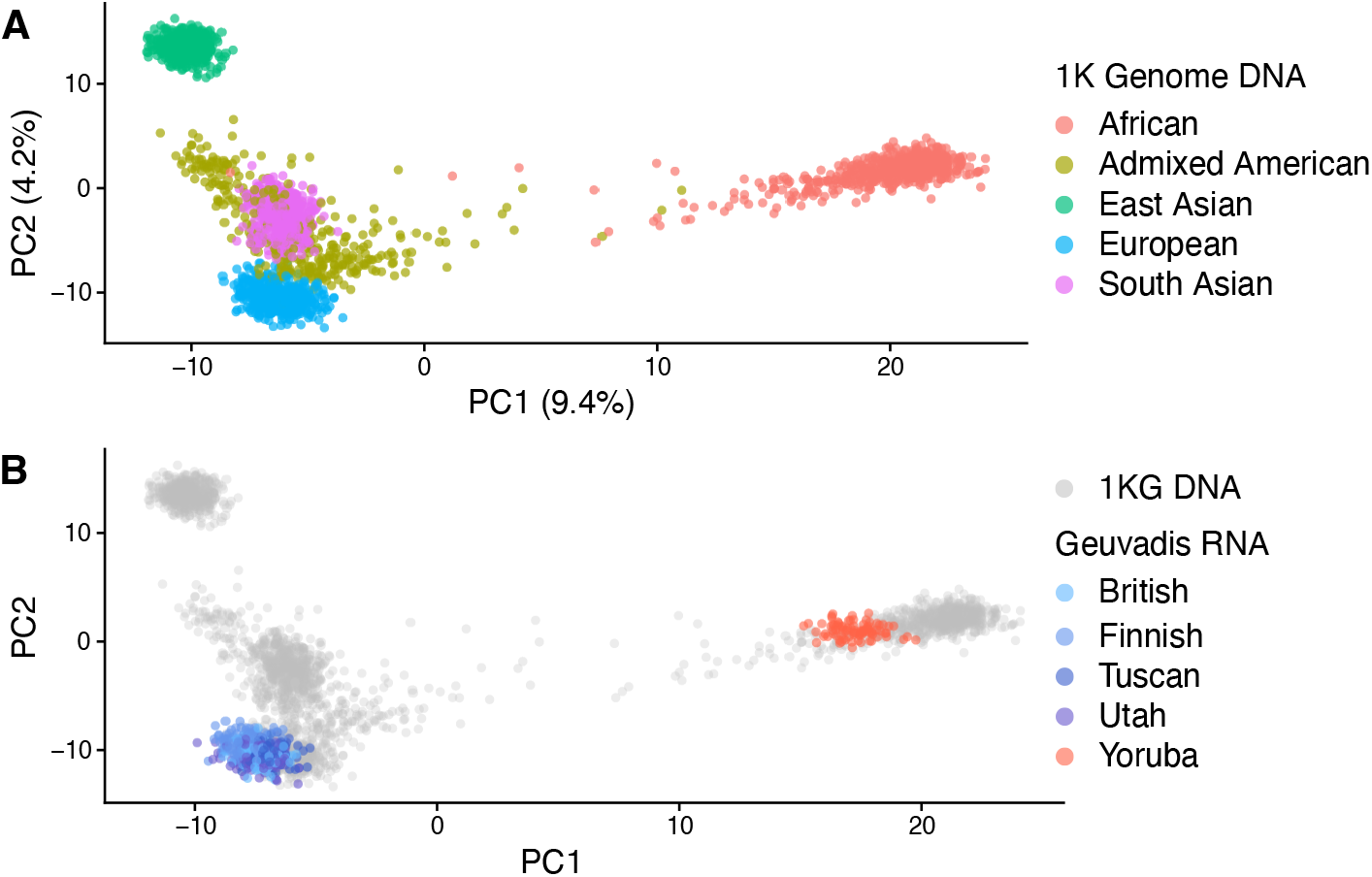
Projection of predicted genotypes onto a reference population space. **(A)** PCA was computed from whole-genome DNA sequencing from the 1000 Genome phase III study and individuals are colored based on the 5 different super-population (East Asian=EAS, South Asian=SAS, African=AFR, European=EUR, American=AMR). **(B)** RNA-seq predicted genotypes from Geuvadis project are projected onto this subspace, and each individual is colored based on 5 annotated populations (British, Finnish, Tuscan, Utah, Yoruba). Grey points are 1000 genomes data.

One of the challenges of analyzing genotypes based on RNA-seq data is the substantial amount of missing genotypes. This is because the expression of many genes is tissue-specific and we can only genotype SNPs in expressed genes. To find SNPs expressed across a wide variety of tissues to use in PCA, we selected 30,875 SNPs which are expressed in a majority of GTEx tissues (Methods). Using these selected SNPs, we first performed PCA on the 1000 genomes data and then projected our predicted genotypes onto this space.

To ensure that our PCA projection is reliable, we projected predicted genotypes from the Geuvadis project onto this reference population subspace. We then compared the location of the Geuvadis samples – based on genotypes from RNA sequencing data – to the location of the 1000 genomes samples – based on the genotypes from DNA sequencing data, and found excellent agreement, validating our approach (Figure 5B).

These promising results encouraged us to extend this analysis to all of SRA. We first restricted our analysis to the 65,844 out of 314,596 (21%) samples from Recount3 SRA samples with equal or less than 10% missing genotype information (Figure 6A). All these samples are placed near known super-populations.

**Figure 6.**
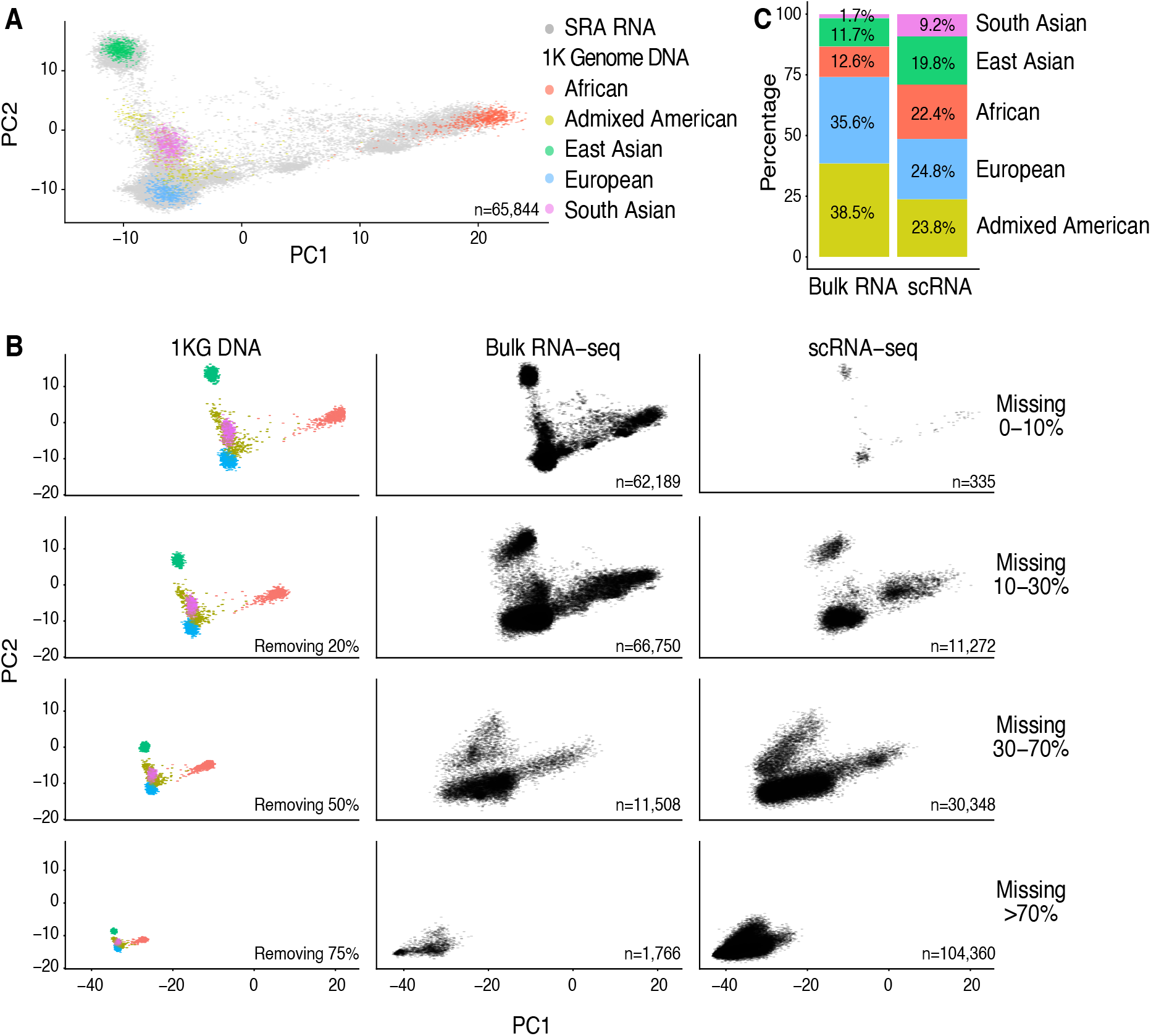
Projection of SRA samples onto 1000 Genomes Project first two principal components. Color dots represent individuals in 1000 Genome study with their corresponding ancestry origin. **(A)** Gray points represent SRA individuals with unknown ancestry information with less than 10% missing predicted genotypes. **(B)** Impact of missingness; First column shows the 1000 Genome super-populations with different proportions of genotypes removed at random. Columns 2 (bulk RNA-sequencing) and 3 (single-cell RNA-sequencing) depict SRA data with different ranges of proportions of missing genotypes. **(C)** Ancestry prediction of SRA bulk and single cell RNA-seq samples. Only samples with an estimated prediction accuracy higher than 70% are depicted.

However, despite selecting SNPs which are broadly expressed across GTEx, we still found that a majority of samples had more than 10% missing genotypes. We encode present genotypes as 1,2,3 with a missing genotype being encode as -1. To examine the impact of this encoding, we randomly removed genotypes from our reference 1000 genomes populations (Figure 6B). Depending on the amount of missing genotypes, the position of the projected points is shifted to the lower left of the display (as expected based on our choice of encoding).

SRA contains a roughly equal proportion of bulk and single-cell RNA-sequencing runs. When we separate sequencing runs by bulk and single-cell, we observe that the single-cell data has a substantial degree of missing genotypes as expected; 71% of the scRNA-seq data had more than 70% missing genotypes. Furthermore, the projected points are shifted as expected (Figure 6B).

Based on this, we developed a method to predict super-population. We used K-means clustering on PCA projections combined with the 1000 Genome reference data as a training set (Methods). We grouped samples within 1% increments of missing genotype to perform prediction (Supplementary Figure S7 and Figure S8). We trained our model by randomly removing genotypes from our training data, followed by projection. We used a threshold of greater than 70% estimated prediction accuracy to indicate a successful prediction. With this definition, we were able to successfully predict the super-population of a majority (121,698 sample, 86%) of bulk RNA-seq and (113,844 samples, 78%) scRNA-seq samples (Figure 6C).

Amongst the bulk RNA-seq studies, Admixed American and European super-populations were both highly represented with 38% and 35.9% of the samples respectively. The next largest super-populations were Africans with 12.6% of SRA samples and East Asians with 11.7%. In contrast, for single-cell RNA-seq studies, we observe a more even spread amongst these four super-populations (23.8%, 24.8%, 22.4%, and 19.8%) and a strong relative increase in the proportion of South Asians (1.7% to 9.2%). The proportions from single-cell RNA-seq studies reflects cells rather than individuals. Nevertheless, this suggests a remaining lack of diversity in genomic data despite the recent efforts.

## Discussion

We have expanded the Recount3 expression repository by providing genotype information on 336,463 samples from SRA and GTEx. We have successfully created a single-sample pipeline that predicts the genotype at biallelic SNPs in expressed regions. This information enables future large-scale genomic analyses, including eQTLs, allele-specific expression, and variant prioritization. Our accurate and efficient genotyper for RNA-seq data provides an opportunity to follow up existing transcriptomic studies with genetic data without additional sequencing. This would not replace gold standard SNP calling technologies such as whole-genome sequencing or array-based genotyping but as an alternative with no additional cost, when RNA-sequencing is already available.

A limitation of our data processing is that we only provide predicted genotypes for SNPs that are present in individuals in the GTEx study. This limitation arises because we need a gold standard set of genotypes to estimate prediction accuracy. It would be easy to deploy and test our model if gold standard data were available on different SNPs. Because of the structure and trainign of our model, we expect a similar performance on a new set of SNPs as we here report on the variants in GTEx. Furthermore, we only predict genotypes for SNPs in coding regions. To expand the number of genotyped SNPs, one could perform whole genome imputation using our predicted genotypes (Das et al., 2016). This would expand our predictions to include non-coding regions.

Our analysis of the composition of the Short Read Archive at the super-population level, shows that Admixed Americans and Europeans are the two largest super-populations. This reinforces the need to further diversify genomic data.

## Methods

### Data

#### Recount3

Recount3 consists of 347,093 human samples aggregated from the unrestricted access part of the Sequence Read Archive (SRA), Genotype-Tissue Expression (GTEx version 8) and The Cancer Genome Atlas (TCGA). All RNA-seq samples have been uniformly processed with the Recount3 Monorail pipeline (Wilks et al., 2021).

Processed RNA-seq samples from Recount3 are stored in the form of total base read count (bigWig files) and alternative read counts (zst files). The total base read count records the total number of reads aligned to the reference genome (GRCh38) at the nucleotide level, and the alternative read counts record the coverage of each alternative nucleotides compared to the reference genome. Read alignment information, such as in the form of BAM files, are not kept in the Recount3 pipeline. The total base read count files are accessible from the Recount3 R package, however, the alternative read count files are not publicly available due to privacy concerns.

We use two datasets within Recount3 for training and evaluating our genotyping and accuracy models:

- The Genotype-Tissue Expression project (GTEx version 8) is used for model training and evaluation. The project consists of 19,081 samples from 972 individuals and 54 tissues. 838 individuals have whole genome DNA sequencing (WGS) performed which serve as genotyping ground truth. We used these 838 individuals for our model training and evaluation.
- The Geuvadis project is used as an out-of-study evaluation set. The project has imputed SNP arrays on lymphoblastoid cell lines for 462 HapMap individuals from 4 European populations and 1 African population.

#### 1k genomes DNA sequencing

The 1000 Genome Phase III study includes 2,504 individuals from 26 different sub-populations and 5 different super-populations, and their Whole Genome Sequence VCF files were downloaded from http://ftp.1000genomes.ebi.ac.uk/vol1/ftp/release/20130502 (T1GP Consortium et al., 2015). The VCF files were lifted over from hg19 to hg38 by using Picard LiftoverVcf and the UCSC reference genome (DePristo et al., 2011).

#### Selection of SNPs for training and genotyping

We only genotype samples at known SNPs in the GTEx population. Using the WGS Variant Call Format (VCF) of 838 individuals, we first selected biallelic SNPs using bcftools version 1.2 (Danecek et al., 2021). We further filtered our SNPs to protein coding regions using Ensembl version 85 for GRCh 38 reference genome. Finally, we remove SNPs belonging to sex chromosomes as we assume that both alleles are expressed unless in the extreme case of imprinting. This gives us 20,980,266 SNP locations.

### Data transformations

#### Computing coverage of reference and alternative allele

The coverage of the alternative allele at the nucleotide level is computed by summing all possible alternative alleles’ read counts via the alternative read count file. For instance, if the reference allele is “A”, then the coverage of the alternative allele is the sum of coverage for “T”, “C”, and “G” alleles. Then, the coverage of the reference allele at the nucleotide level is computed by subtracting the alternative allele coverage from the total base coverage via the total base read count file.

#### *M* and *S* transformation

Let ref_*i,j*_ be the reference counts for SNP *i* from sample *j*, and alt_*i,j*_ be the alternative counts for SNP *i* from sample *j*. For model training, multiple samples *j =* 1, …, *J* of the same tissue type are analyzed together. For genotyping, only one sample *j =*1 is analyzed at once so the *j* index is dropped. We transform ref_*i,j*_ and alt_*i,j*_ into *M*_*i,j*_ and *S*_*i,j*_ values with a pseudocount of 1, based on (Carvalho et al., 2010):

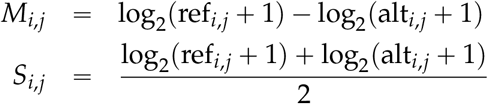

### Genotyping Model

We developed a model to genotype RNA-seq samples using reference and alternative read counts available through the Recount3 project, using *M* and *S* values (Data transformations).

#### Model overview

Let *Z*_*i,j*_ ∈{AA, AB, BB} be the 3 possible genotypes for SNP *i =*1, …, *I* from sample *j=* 1, …, *J* representing reference homozygous, heterozygous, and alternative homozygous alleles, respectively. The genotype is treated as a discrete latent variable. As prior we use a multinomial distribution:

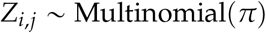

with

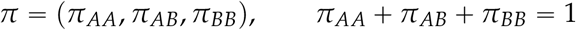

Then, conditional on genotype *Z*_*i,j*_ *= g* with *g* ∈*{*AA, AB, BB}, we assume that the relationship between *M*_*i,j*_ and *S*_*i,j*_ can be described by a mean function *f*_*g*_(*s*) and variance function 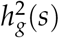:

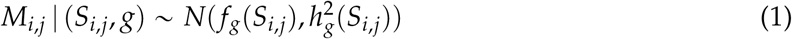

We assume a linear relationship between *M*_*i,j*_ and *S*_*i,j*_ to parameterize *f*_*g*_(*S*_*i,j*_), and a smooth function between variance of *M*_*i,j*_ and *S*_*i,j*_ to parameterize 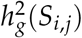.

Given a sample to be genotyped with values of *M*_*i*_, *S*_*i*_ (index *j* is dropped due to *J =*1), we use model parameter estimates of 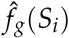 and *ĝ*_*g*_(*S*_*i*_), and prior *π* to compute the posterior probability for each possible genotype *Z*_*i*_ *= g* ∈*{*AA, AB, BB}:

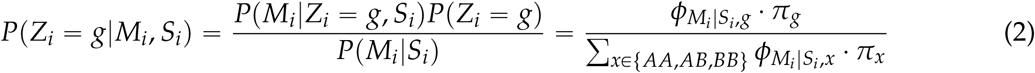

where 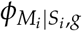 is the normal likelihood evaluated at *M*_*i*_ with mean and variance described in Equation 1:

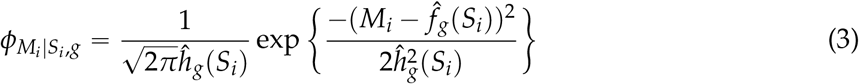

We use the maximum a posteriori probability (MAP) estimate of the posterior to predict the genotype.

#### Construction of GTEx training and testing set

To build our training set, we wanted to select a wide variety of tissues from GTEx so that the genotyping and accuracy models will be robust to a wide range of tissues. We first selected 33 tissues such that each tissue has at least 200 individuals profiled for RNA-seq and WGS. The union of individuals in the 33 tissues encompassed all 838 individuals in GTEx with RNA-seq and WGS. We then randomly sampled 638 individuals for the training set, and for each individual in the training set, we sampled one tissue sample for the training set. Thus, we have 638 samples from 638 individuals for the training set, with equal contribution from every individual.

The remaining 200 individuals with RNA-seq and WGS in GTEx are then designated for the test set. Here, each individual may have multiple samples from multiple tissues. We get a total of 3,901 samples from 200 individuals in the test set.

#### Model training

We used our GTEx training set of 638 samples from 638 individuals for model training. Starting with our candidate set of SNPs, we removed SNPs with an average coverage across samples less than 3. Then, we transformed our reference and alternative counts into *M*_*i,j*_ and *S*_*i,j*_ values.

##### Averaging *M* and *S* values

We noticed that within a genotype, the correlation between averaged *M* value for a SNP across samples with averaged *S* value for a SNP across samples is higher than the correlation between *M* and *S*. We therefore use the mean of *M*, the mean of *S*, and the standard deviation of *M* across samples with a given genotype in the training set as a more robust way to describe relationship between *M* and *S*. This yields mean(*M)*_*i,g*_, mean(*S)*_*i,g*_ and sdp*M*q_*i,g*_ with *i =* 1, …, *I, g* ∈ {*AA, AB, BB*}. As a short-hand, below we use 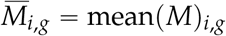 and 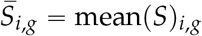

##### Parameter estimates of 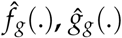, and *π*

For each genotype *g*, we fit a mean model with mean(M) as the response variable and mean(S) as the predictor and a variance model with sd(M) as the response and mean(S) as the predictor, giving us a total of 6 models.

The mean model for genotype *g* is described as:

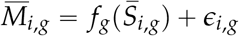

where 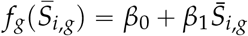 and *ϵ*_*i,g*_ is a mean-zero normal variable.

The variance model for genotype *g* is

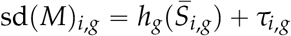

where 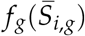 is a generalized additive model (GAM) where sd (*M)* _*i,g*_ is the outcome and 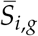 is the predictor, and *τ*_*i,g*_ is a mean-zero normal variable.

Lastly, the prior *π =(π*_*AA*_, *π*_*AB*_, *π*_*BB*_) is estimated as the proportion of SNPs from all samples that have the genotype in the GTEx training set across all tissues.

#### Genotype prediction

Given a new sample from Recount3, we select the subset of candidate SNP locations that have a coverage greater than 4. For each SNP, we compute the *M*_*i*_ and *S*_*i*_ transformation. Then we compute the normal likelihood, 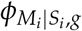, evaluated at *M*_*i*_ for each genotype *g* as described in Equation 3. Finally, we calculate the posterior probabilities for *Z*_*i*_*= g* ∈*{AA, AB, BB}* based on 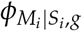 and prior *π* via Equation 2. We use the MAP estimate as our genotype prediction.

We show our genotype prediction boundaries in Figures 1b and 2C. We noticed that in our decision boundary there is a potential region for misclassification around *S* 8 and *M* 4: this region is predicted to be “BB” alternative homozygous, when clearly it should be “AA” reference homozygous or “AB” heterozygous. This is due to the likelihood function for “BB” slightly dominating the likelihood function of “AA” and “AB”. We checked our evaluation data to see how many SNPs is misclassified in this region. In the GTEx test set and Geuvadis out-of-study set, the probability of this misclassification is on the order of 10^− 6^, so this is not a concern.

#### Model evaluation

We evaluated our genotype model on the GTEx test set consisting of 3901 samples from 200 individuals and the Geuvadis out of study set consisting of 462 samples and individuals. We report our genotyping model accuracy for each SNP from all samples, using genotypes from WGS or array-based technologies as the gold-standard. The accuracy is calculated at the allelic level, which implies that each biallelic SNP can have an accuracy of 0/2 = 0, 1 /2 = 0.5, or 2/2 = 1. We have three definitions of allelic accuracy: “Overall Accuracy” considers the allelic accuracy of both alleles, “Major Accuracy” considers the allelic accuracy of the major allele, and “Minor Accuracy” considers the allelic accuracy of the minor allele. The major allele for a SNP is defined as the most prevalent allele for the SNP across the 838 GTEx individuals computed using gold-standard WGS, and the minor allele is the least prevalent allele. We use the following rule to compute our three definitions of allelic accuracy, assuming “A” as major allele and “B” as the minor allele in Table 1.

We report the tissue’s or study’s accuracy as the average of each SNP’s accuracy, after removing all NA values.

### Accuracy Model

We developed an accuracy model to predict the accuracy of our genotyping model when the gold-standard genotype is unknown. For this model, we compute genotyping accuracy at the genotype level so that each SNP from a sample have an accuracy of 0 or 1.

#### Model overview

Denote Acc_*i,j*_ ∈{0, 1} to be the genotype accuracy for SNP *i=* 1, …, *I* from sample *j=* 1, …, *J*. For model training, multiple samples *j =*1, …, *J* of multiple tissue types are analyzed together. For predicting the accuracy, only one sample *j =*1 is analyzed at once so the *j* index is dropped. Also, denote coverage of SNP *i* from sample *j* as Cov_*i,j*_. Lastly, the major allele frequency for a SNP is defined as the number of most prevalent allele for the SNP across the 838 GTEx individuals divided by the number of total alleles for the SNP across individuals. This is computed using the gold-standard WGS. We create categorical variable 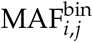 based on which bin the major allele frequency belongs to: [0.5, 0.632), (0.632, 0.749], (0.749, 0.84], (0.84, 0.908], (0.908, 0.95], and [0.95, 1]. The bins between 0.5 and 0.95 are determined from equal sized quantiles of major allele frequencies from the GTEx training set.

We observed that the majority of SNPs have a major allele frequency belonging to the 0.95, 1 bin (Supplementary Figure S3) and the relationship between *Acc*_*i,j*_ and *Cov*_*i,j*_ is different than other bins (Supplementary Figure S5). Therefore, we separate our training SNPs into two groups based on whether their major allele frequency bin is [0.95, 1] or not. Each SNP group is modeled separately but with the same parameterization as described below.

We modeled the genotype accuracy as a function of coverage and major allele frequency bin via a logistic regression model (written using model formulas):

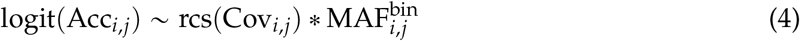

where rcs(·) is a restricted cubic spline function with knots at coverage values of 4, 6, 10, 16, and 40.

#### Model training

We used our GTEx training set of 638 samples from 638 individuals for model training. Starting with our candidate set of SNPs, we removed SNPs with an average coverage across samples less than 4. We separate our training SNPs into two groups based on whether their major allele frequency bin is [0.95, 1] or not and we fit two accuracy models with the same parameterization as described in Equation 4.

#### Model prediction and evaluation

We evaluated our accuracy model on the GTEx test set consisting of 4901 samples from 200 individuals and the Geuvadis out of study set consisting of 462 samples and individuals. For each sample and our candidate set of SNPs, we kept SNPs with coverage greater than 4.

Once we predict the genotyping accuracy for SNPs from the test set samples, we group SNPs together that have the same coverage and major allele frequency bin to compute the predicted mean genotype accuracy, which is a continuous value between 0 and 1. Similarly, we compute the ground truth mean genotype accuracy. We examine the model fit by comparing the mean prediction accuracy and ground truth mean genotype accuracy graphically (Figure 4, Supplementary Figure S6), and quantitatively compute the absolute value difference between these two values.

#### Population structure analysis

We obtained a VCF file for the 1k genome data (See the Data section). Genotype values were mapped to values of 1, 2, and 3 for reference homozygous, heterozygous, and alternative homozygous respectively before PCA was performed. To select the SNP locations for PCA analysis, we choose 30,875 SNPs where we had genotype prediction for all the samples in our GTEx evaluation tissues with more than 80 samples.

To project our predicted genotypes onto the subspace, we first selected the same SNP locations. Missing genotype information (due to insufficient coverage) were given the value of -1, and genotyping values were mapped to values of 1, 2, and 3 as before. New samples from Recount3 were projected onto 1000 Genome PCA by using the rotation matrix estimated from PCA of the 1k genome data and the predicted genotypes where standardized using scale and center values calculated using the 1k genome data as well. Calculations were performed using `prcomp()` and `predict()` function from the `stats` R package.

#### SRA population structure prediction

To predict the unknown population structure of SRA samples, we trained a model using K-means clustering method from `caret` R package. To deal with different amounts of missing genotypes in our samples, we grouped samples based on their missing genotype in 1% increments. For each group, we simulated 1000 Genome by assigning random SNPs a value on -1 (missing genotype). We then trained the model with this simulated data by using k=6. Using PC1 and PC2 values as predictors in `predict()` function, we obtained the super-population category for each sample. We also obtained the posterior probability by using the argument `type = “prob”`. We only selected bulk RNA-seq and samples with prediction probability of more than 70% for downstream analysis.

## Data and code availability

We are currently communicating with NIH to finalize the details on data access procedures. Until that is finalized, the genotyping calls are not publicly available. The code used to produce the manuscript is available at https://github.com/hansenlab/recount_genotype_paper. The R package for single sample genotype calling is available at https://github.com/hansenlab/RecountGenotyper.git

## Contributions

AR and CCL contributed equally to conceptualization, method and software development, data analysis and writing. SW contributed to conceptualization, preliminary data analysis and methodology. JTL and KDH contributed conceptualization, supervision, funding acquisition, writing. All authors read ana approved the final version of the manuscript.

## SUPPLEMENTARY MATERIALS

## 1 Supplemental Figures

**Supplementary Figure S1.**
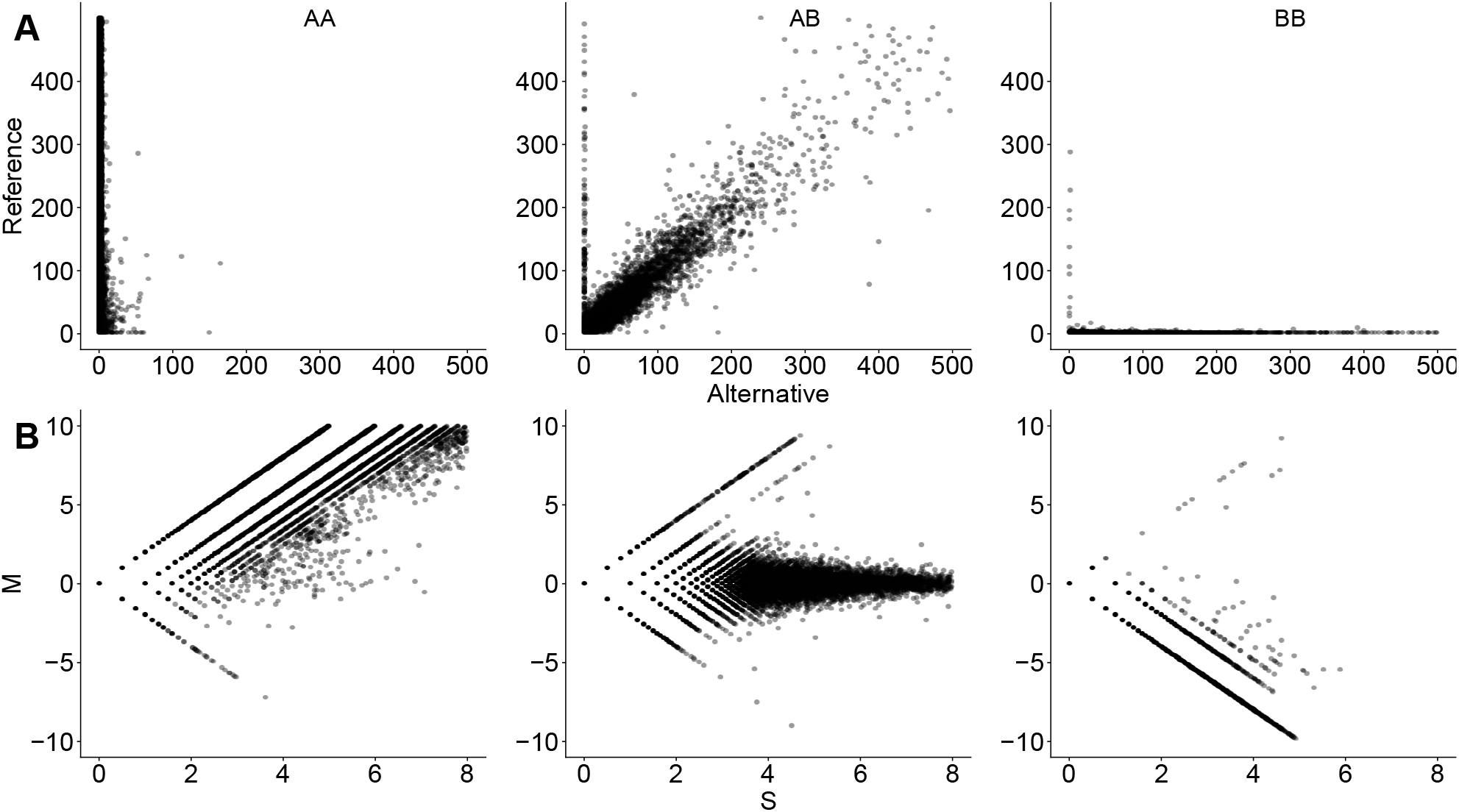
Transformation of SNP coverage values. We depict 200 random GTEx samples in the training set. **(A)** Raw reference and alternative read counts where each point represents a SNP of a sample. Each plot is faceted based on genotype (A being reference and B being alternative). **(B)** The same samples and SNPs are plotted again using *S* vs. *M* transformation.

**Supplementary Figure S2.**
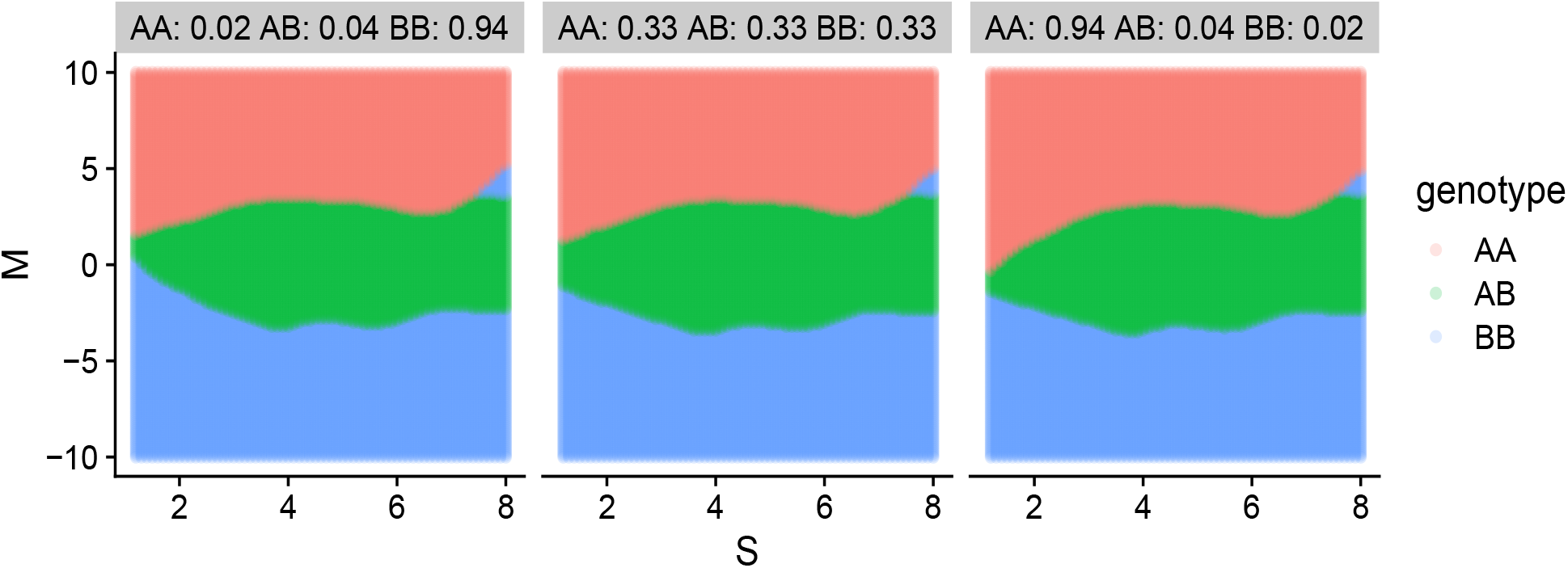
Sensitivity analysis for the choice of prior distribution. Each facet depicts the decision boundaries of the three genotypes based on a prior genotyping distribution. The prior genotyping distribution are specified in each facet’s title. The reference allele is “A” and the alternative allele is “B”.

**Supplementary Figure S3.**
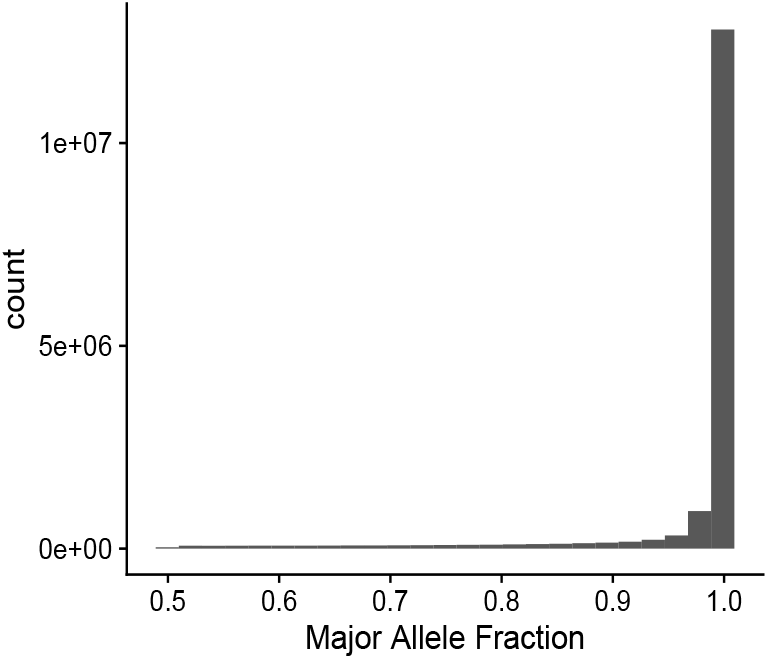
Distribution of major allele frequency of GTEx test set. The major allele frequency for a SNP is the number of most prevalent allele for the SNP across the 200 GTEx test set individuals divided by the number of total alleles for the SNP across the individuals.

**Supplementary Figure S4.**
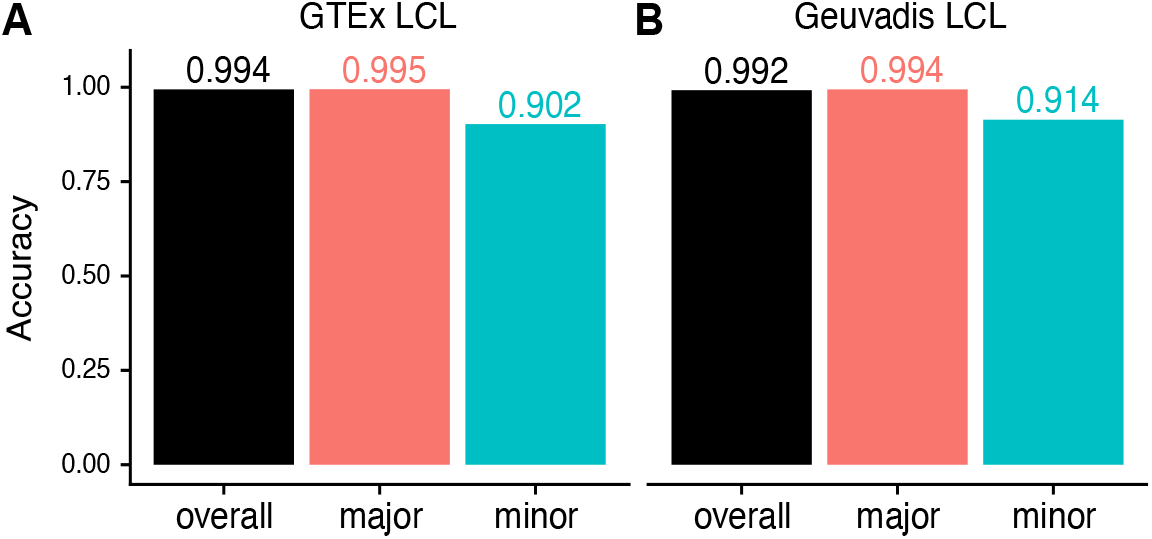
Model performance for lymphoblastoid cell lines. **(A)** Model performance for 35 GTEx LCL samples in the GTEx test set. **(B)** Model performance for 462 Geuvadis samples as an out-of-study set.

**Supplementary Figure S5.**
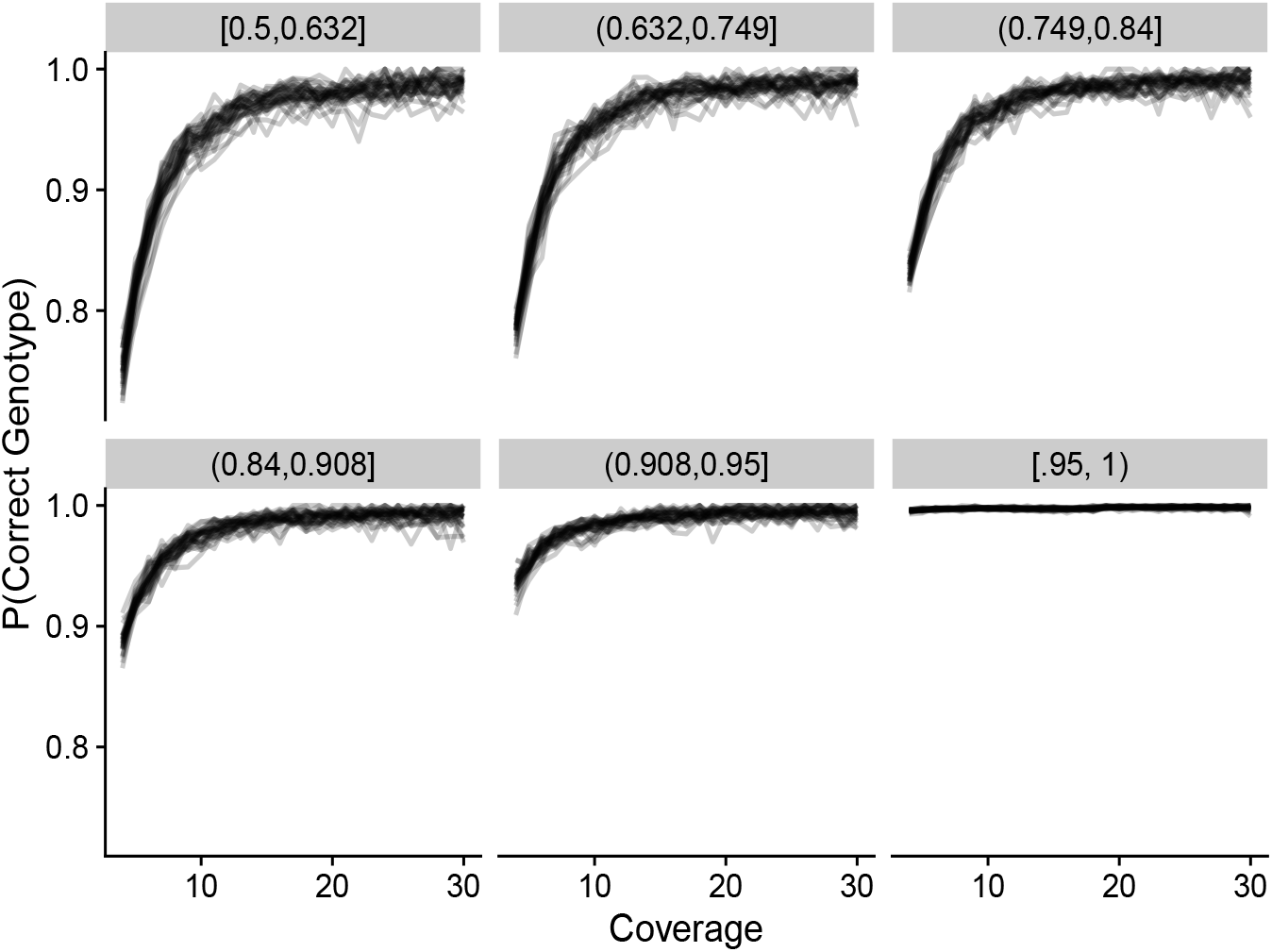
Genotyping accuracy as a function of sequencing coverage and major allele frequency. Genotyping accuracy at the SNP level is grouped by discrete coverage values and major allele frequency bin to get a continuous averaged accuracy value. Each facet denotes a major allele frequency bin, and each line represents a tissue type from GTEx training set.

**Supplementary Figure S6.**
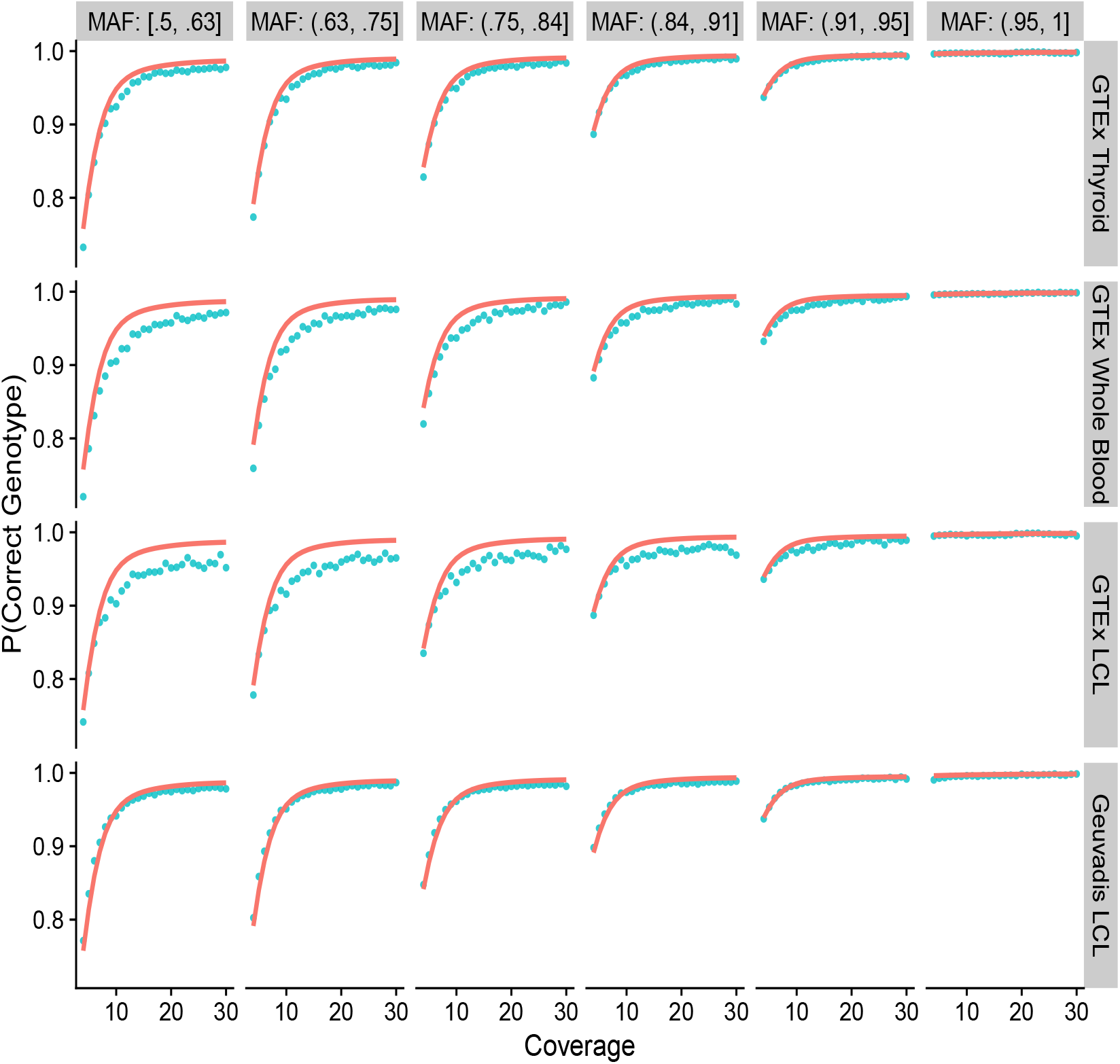
Genotyping accuracy as a function of sequencing coverage and major allele frequency for several tissues. Genotyping accuracy at the SNP level is grouped by discrete coverage values and major allele frequency bin to get a continuous averaged accuracy value (blue-green points). The model prediction is shown in red lines. Each facet along the x-axis denotes a major allele frequency bin. Each facet along the y-axis denotes a different tissue type. We contrast a well fitted tissue type (GTEx Thyroid) against less well fitted tissue types (GTEx Whole Blood, GTEx LCL). We also contrast model predictions of LCL tissues between two studies (GTEx LCL vs. Geuvadis LCL).

**Supplementary Figure S7.**
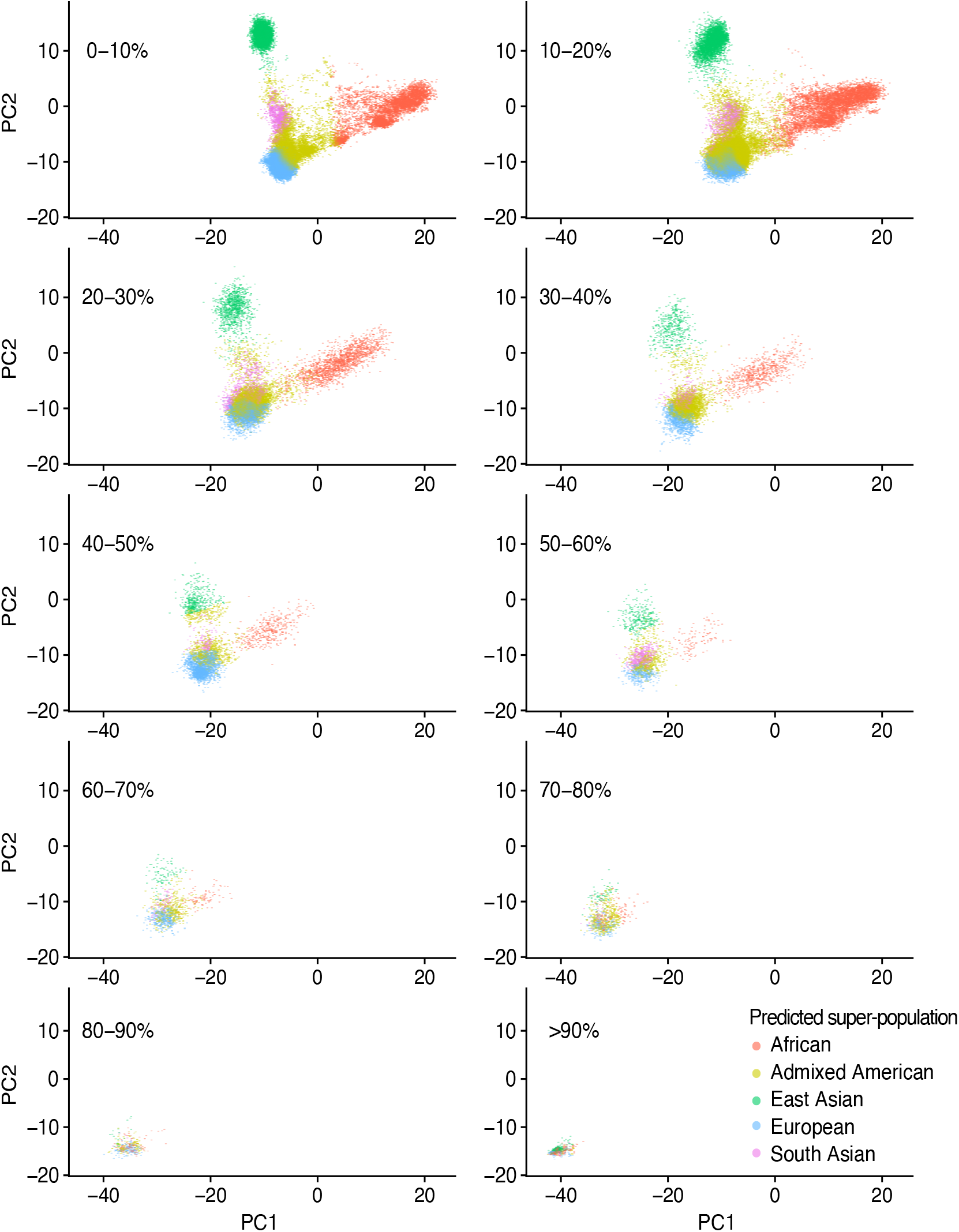
SRA Bulk RNA-seq samples ancestry prediction. Bulk RNA-seq samples from SRA are separated based on their percent missing genotype in 10% increments. Each color corresponds to the predicted super-population based on 1000 Genome reference.

**Supplementary Figure S8.**
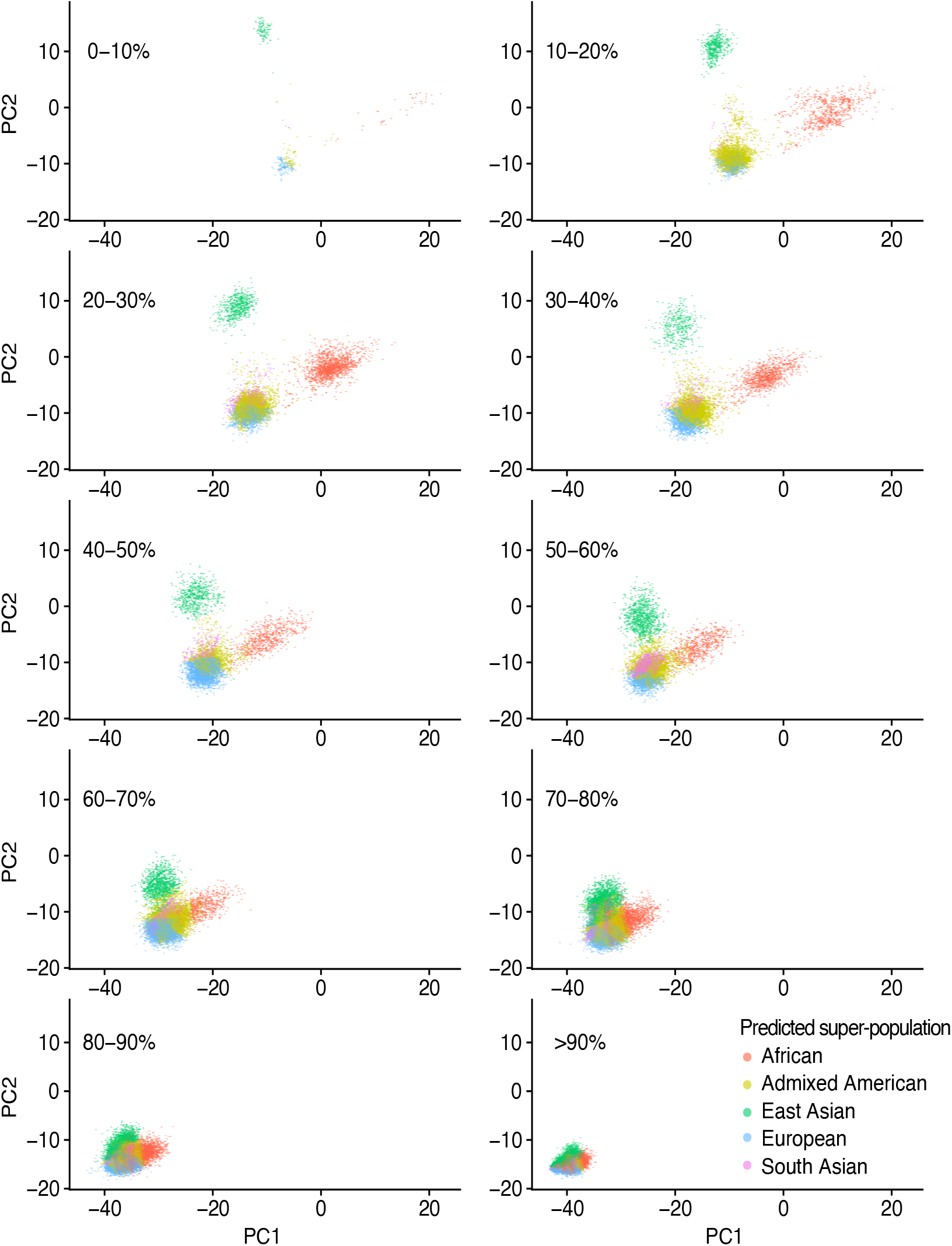
SRA single-cell RNA-seq samples ancestry prediction. Single-cell samples from SRA are separated based on their percent missing genotype in 10% increments. Each color corresponds to the predicted super-population based on 1000 Genome reference.

